# SELINA: Single-cell Assignment using Multiple-Adversarial Domain Adaptation Network with Large-scale References

**DOI:** 10.1101/2022.01.14.476306

**Authors:** Pengfei Ren, Xiaoying Shi, Xin Dong, Zhiguang Yu, Xuanxin Ding, Jin Wang, Liangdong Sun, Yilv Yan, Junjie Hu, Peng Zhang, Qianming Chen, Taiwen Li, Chenfei Wang

## Abstract

The rapid accumulation of single-cell RNA-seq data has provided rich resources to characterize various human cell types. Cell type annotation is the critical step in analyzing single-cell RNA-seq data. However, accurate cell type annotation based on public references is challenging due to the inconsistent annotations, batch effects, and poor characterization of rare cell types. Here, we introduce SELINA (single cELl identity NAvigator), an integrative annotation transferring framework for automatic cell type annotation. SELINA optimizes the annotation for minority cell types by synthetic minority over-sampling, removes batch effects among reference datasets using a multiple-adversarial domain adaptation network (MADA), and fits the query data with reference data using an autoencoder. Finally, SELINA affords a comprehensive and uniform reference atlas with 1.7 million cells covering 230 major human cell types. We demonstrated the robustness and superiority of SELINA in most human tissues compared to existing methods. SELINA provided a one-stop solution for human single-cell RNA-seq data annotation with the potential to extend for other species.

## Introduction

Single-cell RNA sequencing (scRNA-seq) can profile thousands of cells to reveal heterogeneity within complex tissues. The key step in scRNA-seq data processing is cell type annotation, which is vital for interpreting function features for certain cell types and is required for many downstream analyses, including trajectory analysis or cell-cell interactions. Traditional cell type annotation methods can be roughly divided into two categories. Marker gene-based methods including Garnett^1^ and SCINA^2^ rely on clustering and quality of cell type-specific marker genes. Reference data-based methods, such as scmap^3^, scPred^4^, SingleR^5^, CHETAH^6^, SingleCellNet^7^, and ACTINN^8^, transfer cell type labels from a reference dataset and are independent of clustering results and marker genes. With the continuous accumulation and increased throughput of single-cell datasets, reference data-based methods show improved accuracy and broader applications compared to marker gene-based methods.

Although reference data-based methods have the above advantages in cell type annotation, several challenges remain to be resolved. First, current tools are often designed for transferring cell type assignments between single reference data and single query data; hence, they cannot leverage the wealthy information hidden in the enormous public data. Second, the cell numbers of different cell types are often imbalanced; therefore, the minority cell types are always ignored in the modeling process. Third, the underlying batch effects between reference data and query data are often overlooked, which may greatly hinder accurate label transfer. Last, all these methods heavily rely on the quality of reference datasets. However, none of them provide users with comprehensive reference datasets. Even though great efforts in systematically collecting and curating public datasets have been made to build scRNA-seq data portals that involve millions of cells, which spawns Human Cell Atlas (HCA)^9^, Animal Cell Atlas^10^ (ACA), Single Cell Portal from the Broad Institute^11^, Human Cell Landscape^12^ (HCL) and Single Cell Expression Atlas from European Bioinformatics Institute^13^ (EMBL-EBL), there still lacks a uniform and comprehensive reference atlas due to the inconsistent annotation and large batch effects between datasets.

To address these challenges, we built a comprehensive single-cell transcriptomics data atlas consisting of 136 datasets from 35 human tissues ranging from 7 differrent platforms. Based on the 1,706,710 uniformly processed cells covering 230 cell types, we propose an algorithm that can effectively utilize multiple datasets for single-cell assignment. It applies the synthetic minority oversampling technique^14^ (SMOTE) to boost the number of rare cell types and employs multi-adversarial domain adaptation^15^ (MADA) to update the parameters of the supervised deep learning framework in the pretraining stage. Furthermore, it utilizes an autoencoder to adaptively adjust the pretrained parameters based on the distribution of query data. We demonstrated the power of SELINA in batch removal and systematically evaluated the performance of SELINA with existing tools on 92 datasets from 17 tissues. The comprehensive cell type reference of SELINA and its superior ability in transferring annotation pave the way for users to accurately annotate single cells.

## Results

### Overview of SELINA

The SELINA workflow is composed of two steps, reference construction and cell type prediction. For reference construction, public scRNA-seq data were collected from multiple databases and processed with a standardized pipeline from MAESTRO^16^ including quality control, principal component analysis (PCA), batch removal within a dataset, unsupervised clustering, cell type-specific marker evaluation, and annotation (Supplementary Fig. 1a). Next, the inconsistent annotations across datasets are manually unified with the knowledge that the corresponding papers provide (Supplementary Fig. 1b). Finally, the curated cell types are organized into a cell type ontology tree.

Based on the uniformed reference data, a cell type prediction algorithm consisting of three steps is developed. Usually, the classifier will reach a higher training accuracy at the cost of the minority class being misclassified; thus, to increase the sensitivity of classifiers to the minority class, SELINA utilizes SMOTE^14^ to generate synthetic samples so that the classifier can pay more attention to the minority cell types. Then, SELINA takes datasets from one tissue as input and employs a MADA^15^ network to obtain a pre-trained model. By training the supervised deep learning framework in an adversarial way, the underlying common information of the same cells from different batches is uncovered. To further remove the batch noise between the reference data and query data, an autoencoder is used to adaptively fine-tune the pre-trained parameters according to the distribution of query data. Finally, the labels from reference datasets are transferred to the query data based on the fully trained model (Fig. 1).

**Figure 1.**
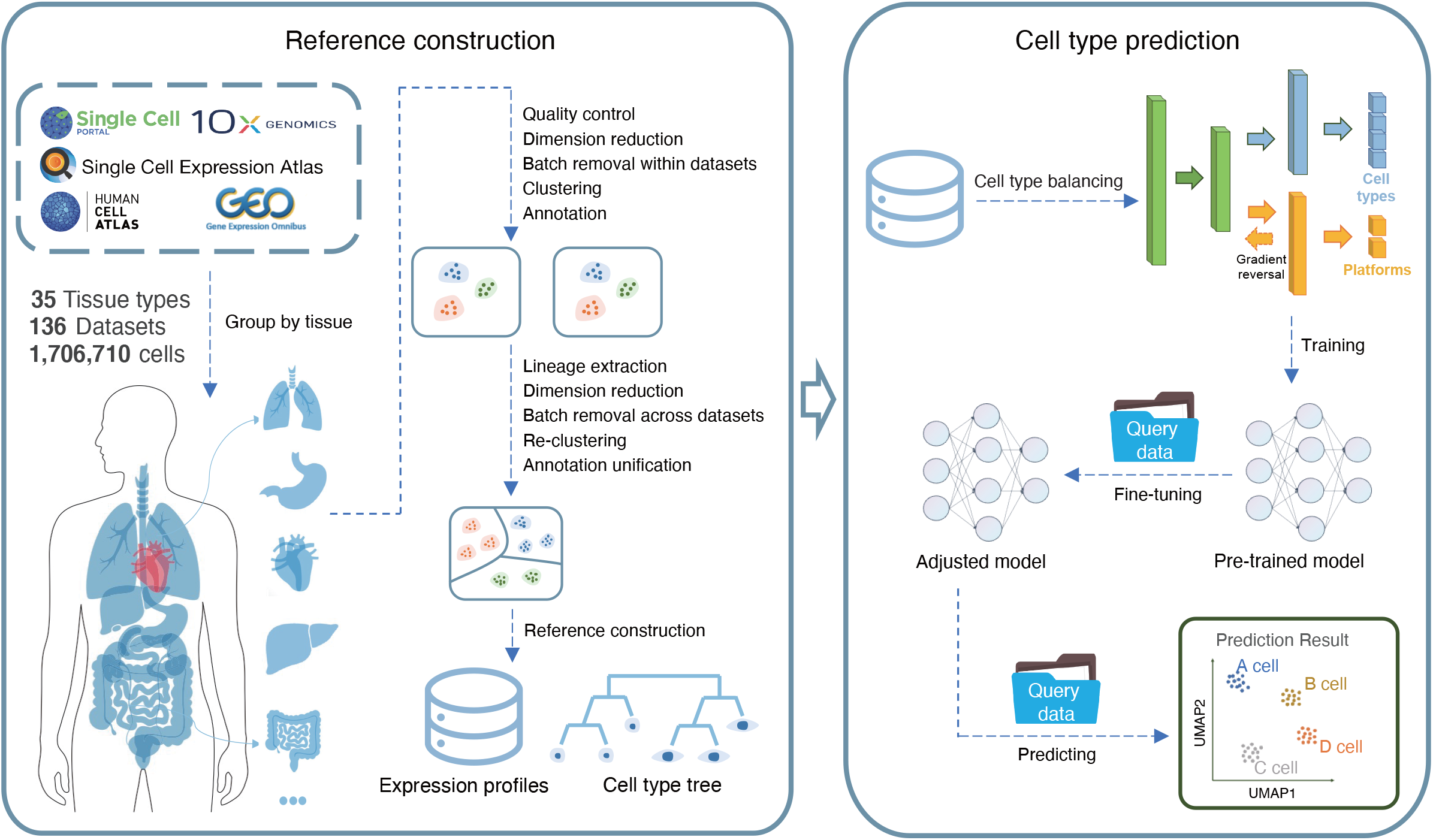
Overview of SELINA. SELINA consists of two sections: reference construction and cell type prediction. Public data were collected from various databases and uniformly processed based on a two-step approach containing within datasets processing and across datasets unification (Methods). For each tissue, a well-organized cell type classification tree was built, and the expression profiles were merged as the algorithm input. The algorithm consists of three steps: cell type balancing, training and fine-tuning. First, the rare cell types from the merged training data are oversampled. Second, the training data is pretrained with a supervised deep learning framework containing a gradient reversal layer. Third, the parameters of the pretrained model are adjusted according to the distribution of query data. Finally, the adjusted model takes query data as input and assigns the cells with cell types from reference data.

### SELINA provides rich sources of single-cell expression profiles with a well-organized cell type ontology tree

In the reference construction step, a total of 1,706,710 cells from 136 datasets were collected to build a single-cell transcriptomic data portal that covered 230 human cell types and 7 different sequencing platforms (Supplementary Table 1). The reference atlas was expanded on the basis of HCL^12^ to include more datasets from other sequencing platforms. All the datasets were categorized into 35 major tissues according to the definition in HCL. We first summarize the dataset features in the SELINA reference. The blood (n=14), intestine (n=14), and bone marrow (n=13) tissue have the richest dataset number (Fig. 2a). Blood and bone marrow tissue also have the largest number of cells, indicating a better characterization of immune cells for our reference. Importantly, 27 out of the 35 tissues have two or more datasets in the reference, suggesting the universal coverage and depth of our reference (Fig. 2a). For each tissue, the inconsistent cell type names between different datasets were first unified and subsequently divided into major lineages and minor lineages based on the literature and Cell Ontology^17^. Taking the cell type names from the liver as an example, the major lineage has 14 different cell type categories, and only dendritic cells (DCs), endothelial cells and epithelial cells have minor lineages (Fig. 2b).

**Figure 2.**
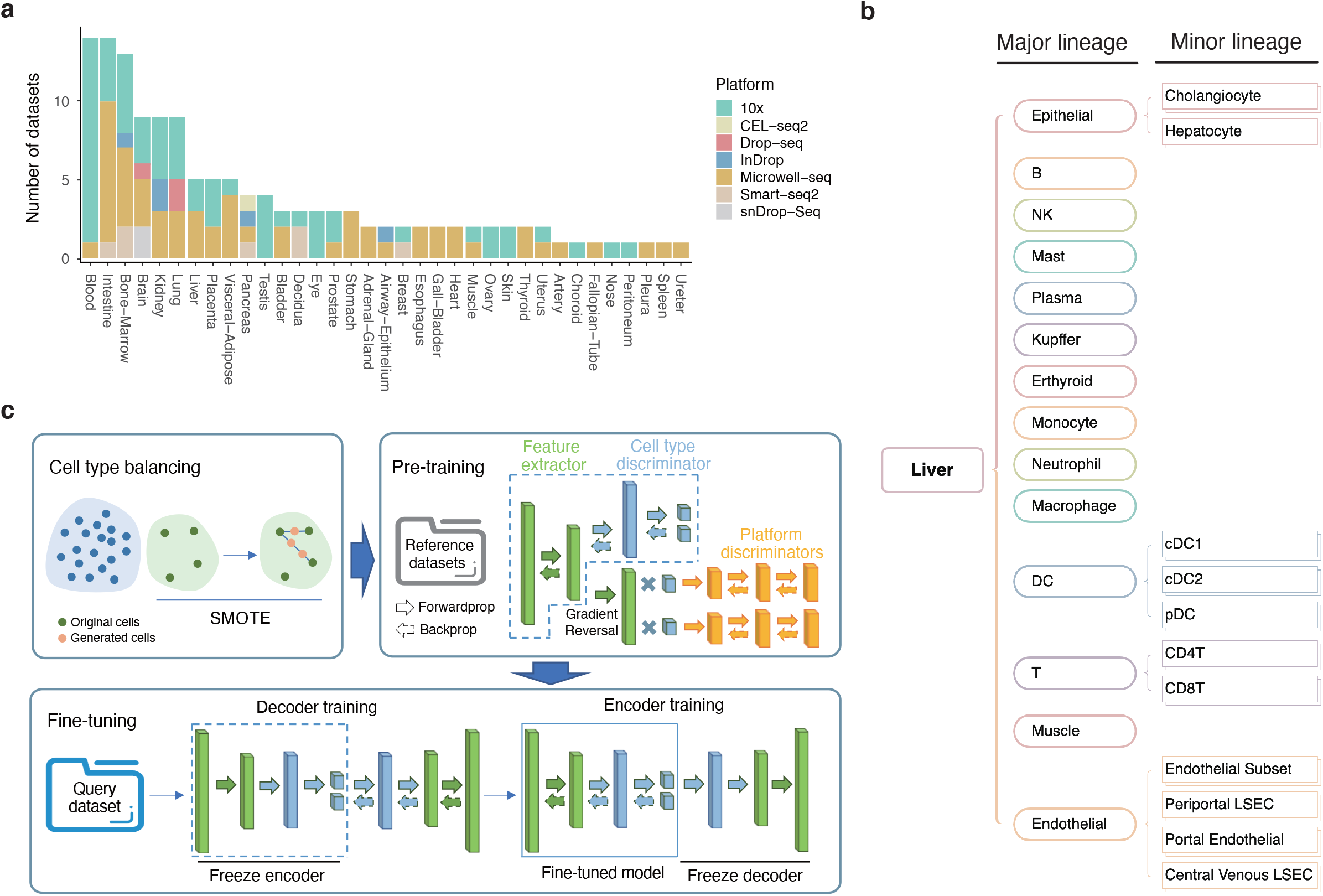
Reference data and algorithm of SELINA. **a.** Datasets number of different tissues collected in SELINA reference. Different colors represent different sequencing platforms. **b.** Cell type classification tree from liver with two levels of annotation. **c.** Annotation algorithm of SELINA. First, the rare cell types are oversampled with SMOTE. The green dots are cells from the original minority cell types, and the orange dots are the synthetic cells. The balanced data are fed into the pretraining framework, which is trained using MADA. Then the pretrained model is fine-tuned using an autoencoder. The feature extractor and cell type discriminator are extracted to construct the encoder, and the decoder is randomly initialized with the structure symmetrical to that of the encoder. The decoder is trained with the encoder fixed; similarly, the encoder is trained with the decoder fixed. Finally, the decoder is removed, and the encoder is used to classify query cells.

Aggregating all tissues together, we constructed a comprehensive human cell type ontology tree based on scRNA-seq data. Previous Cell Ontology has presented hierarchical relationships between 2331 cell types. However, only a small fraction of these cell types can be characterized by scRNA-seq due to experimental capture issues and limited cell numbers in each dataset. In addition, the terminology used to describe cell types in published single-cell studies does not correspond well with the cell type names from Cell Ontology. Therefore, the cell type tree in SELINA only defines the parent-child relationship of cell types that are present in scRNA-seq data. By incorporating and curating cell types from previous studies, SELINA affords users a more unified standard that dictates cell type landscapes in the scRNA-seq data and organized this landscape for annotating input scRNA-seq data.

### SELINA combines MADA and an autoencoder to transfer labels between reference data and query data

The annotation algorithm in SELINA comprises three steps, including cell type balancing, pretraining, and fine-tuning (Fig. 2c). SELINA first adopts SMOTE to oversample the minority cell types. For each cell in a pair of randomly selected cells from rare cell types, SELINA multiplies its gene expression vector by a random weight and then sums the pair of weighted vectors to obtain a synthetic cell which is at a random point on the line connecting the pair of cells. Colloquially, the generated cell is the linear combination of original cells. This process will proceed until the rare cell types reach the same magnitude as the majority cell types or are not less than 1000.

In the pretraining phase, SELINA applies MADA to remove the batch noise caused by different sequencing platforms. The architecture of the pretraining framework consists of three components: a feature extractor, a cell type discriminator and a sequencing platform discriminator. The feature vector generated by the feature extractor will flow into the cell type discriminator and platform discriminator simultaneously. Different from the conventional adversarial neural network^18^, the platform discriminator in our pretraining framework contains multiple classifiers of which the number is equal to the cell types; that is, each classifier is paired with one cell type. For a certain platform classifier, the input feature vector will be multiplied by the probability of the input cell being assigned to the cell type paired with this classifier, and the probability is calculated by the cell type discriminator. During the backward propagation of platform predicting errors, the gradient of the feature vector will be reversed so that the feature extractor is trained to maximize the loss of the domain discriminator; at the same time, the domain discriminator is trained to minimize the loss. Thus, as the training proceeds, features generated by the feature extractor worsen for the platform classifier to classify; however, even though the difference of the input features from different platforms is slight, the platform classifier can always manage to distinguish the platform sources until the difference is nearly eliminated. This strategy can enable fine-grained alignment of expression distributions from different sequencing platforms by capturing the batch information of each cell type separately and training the feature extractor and platform discriminator in an adversarial way.

In the fine-tuning step, an autoencoder is employed to reconstruct the expression profile of query data such that the pretrained model can better fit the query data. The encoder, combining the feature extractor and cell type discriminator from the pretrained model, has learned the transformation from a large amount of reference data. By contrast, the decoder still needs to be adjusted due to the random initialization of the parameters. SELINA first fixes the encoder and updates the parameters of the decoder. Freezing the encoder can prevent the parameters of the encoder from changing considerably so that the learned transformation in the pretrained model can be well preserved. Once the loss decreases to convergence, the decoder can be approximately regarded as the inverse transformation of the encoder, that is, training the encoder alone will not significantly change the parameters. Thus, SELINA will freeze the decoder and turn to update the parameters of the encoder, and to prevent the overfitting, the encoder training only lasts several epochs. The reconstruction loss will be further decreased after encoder training so that the encoder is shifted based on the distribution of query data, which can reduce the batch noise between reference data and query data.

### Data augmentation for rare cell types and batch removal in both data integration and querying processes improve annotation accuracy

To validate the improvement of SMOTE in rare cell type annotation, we selected 4 datasets from the liver as reference data, including 15,859 cells^12,19^, and the remaining dataset was taken as query data^20^. Seven cell types with limited cell numbers that are present in query data are defined as rare cell types, including CD4T (n=98), plasma (n=462), mast (n=12), pDC (n=32), cholangiocyte (n=391), periportal LSEC (n=358) and portal endothelial cells (n=212). We carried out label transfer tests with and without SMOTE and found that implementation of SMOTE greatly increase the number of rare cell types that are correctly assigned, obviously, nearly 1/3 of CD4 T cells were rescued. In addition, half of the mast cells were correctly annotated after using SMOTE, while all of them were misannotated in the original transfer (Fig. 3a). We calculated the F1 of these cell types and found large improvement except for the peripheral LSEC, of which some were probably misannotated in the reference (Fig. 3b). The great improvement indicates that SMOTE can remedy the overfitting caused by the limited number of rare cell types. To confirm that SELINA can eliminate batch effects in reference data, we applied uniform manifold approximation and projection^21^ (UMAP) to display the features transformed by the feature extractor and cell type discriminator. For the liver datasets, after the feature extractor transformation, cells of the same cell type from different platforms are clustered closer compared to the original embedding, suggesting that with the cell type-specific platform discriminators, the feature extractor can uncover the underlying common features among different sequencing platforms for each cell type(Fig. 3c, Supplementary Fig. 3a). With the majority of the batch effects being removed by the feature extractor, we next evaluated the performance of the cell type classifier. Taking CD4+ T-cells, CD8+ T-cells, and NK cells as examples, our cell type classifier showed clear separation of these three cell types, while the results from the conventional PCA transformation followed by Harmony^22^ failed to distinguish them (Fig. 3d). Interestingly, we noticed that the results from cell type classifier showed better integration of CD8T+ cells compared to results from feature extractors (Fig. 3c-d, Supplementary Fig. 3a-b), indicating a further batch correction. To verify whether fine-tuning using autoencoder can remove the batch effects between the reference data and query data, we selected 4 datasets from the liver as reference data^12,19^, and the remaining dataset was taken as query data^20^. For both reference data and query data, we applied UMAP to the features transformed by the cell type classifier from the pre-trained model and the encoder in the fine-tuned model. Clearly, the fine-tuning decreases the distance between reference cells and query cells, indicating that automatic adjustment can fit the pretrained model to the query data (Fig. 3e).

**Figure 3.**
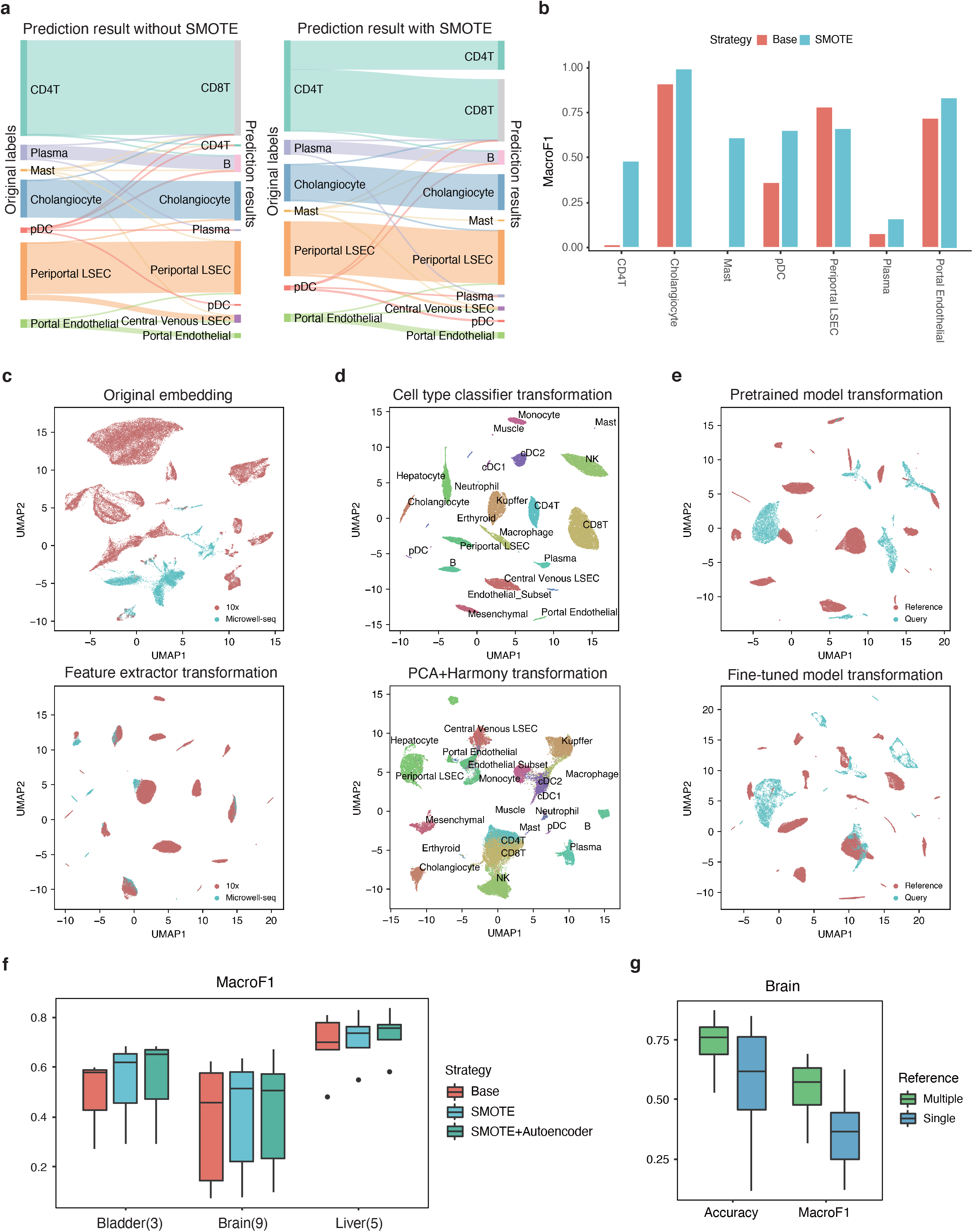
Intrinsic features of SELINA. **a.** The prediction result of rare cell types in query data with and without SMOTE synthesizing new cells in reference data. The height of the bars and linkage lines represents the cell number. The bars on the left are the original labels provided by the corresponding paper, and the bars on the right represent the prediction result of SELINA. **b.** The F1 scores calculated for rare cell types with SMOTE and without SMOTE (Base). **c.** Low-dimensional representation of the reference data from liver before (up) and after (bottom) feature extractor transformation. **d.** Low-dimensional representation of the reference data from liver after cell type classifier transformation and PCA transformation combined with harmony batch removal. **e.** Low-dimensional representation of the query data and reference data before and after fine-tuning. **f.** Mean MacroF1 (3 repeats) of different strategies. Base represents the situation in which the data is not oversampled and the pre-trained model is used without finetuning. SMOTE represents the results when SMOTE is additionally implemented to oversample rare cell types. SMOTE+Autoencoder shows the results when SMOTE and autoencoder are both implemented. **g.** Performance test of multiple datasets and a single dataset used as reference data in brain.

We next investigated whether the performance of SELINA is robust and tested it on three different tissues including bladder, brain, and liver. Each time, we selected one dataset as query data, and the remaining datasets were merged as reference data. MacroF1 is gradually improved with the implementation of SMOTE and autoencoder, whereas the accuracy is relatively stable compared to MacroF1 due to the dominant improvement of rare cell types, which account for a small percentage of the query cells (Fig. 3f, Supplementary Fig. 3c). Moreover, to ensure that effective integration of multiple references can improve the performance, we compared the performance of SELINA using single and multiple reference datasets in the brain and lung tissue, respectively. For single reference tests, each dataset was taken as query data and each of the rest datasets was taken as reference data iteratively. The multiple tests were carried out in the same way as the validation of the strategies used in SELINA. As expected, the performance significantly improves when using multiple reference datasets (Fig. 3g, Supplementary Fig. 3d). Taken together, these results demonstrate that the features implemented in SELINA are robust in different tissues, and an integrated reference will greatly enhance the performance of SELINA.

### SELINA outperforms other existing tools in the comprehensive performance evaluation

To systematically evaluate the performance of SELINA and other existing annotation tools, we selected 92 datasets from 17 tissues covering 185 cell types and 814,788 cells (Supplementary Table 2). For each tissue, we took one dataset as a query dataset each time, and the remaining datasets were merged into one reference dataset; thus, for one tissue, we carried out multiple tests and obtained multiple accuracies and MacroF1. We calculated the average accuracy and MacroF1 and took the highest value in each tissue as 1, and the remaining values from other tools were scaled according to the ratio to the highest value. In terms of the scaled accuracy, SELINA ranks first with an average of 97.51%, followed by ACTINN (92.04%), SingleCellNet (91.48%), mtSC^23^ (89.56%), scibet^24^ (85.98%), singleR (81.52%), CelliD^25^ (78.72%), and scmap (77.17%). For the scaled MacroF1, SELINA also ranks first with an average of 0.9874, followed by mtSC (0.9267), ACTINN (0.9018), SingleCellNet (0.8690), scibet (0.8622), CelliD (0.8478), SingleR (0.8031), and scmap (0.7246). Specifically, among the 17 tissues, the prediction accuracy of SELINA ranked first in 11 tissues and second in 3 tissues, and in terms of MacroF1, SELINA ranked first in 12 tissues and second in 4 tissues (Fig. 4a-b). The detailed performance comparisons for 5 representative tissues are shown in Supplementary Fig. 4a-e. All these results suggest the robustness and high accuracy of SELINA in annotating unlabeled datasets. In addition, we found that compared to the traditional strategies, the deep learning-based model (SELINA, mtSC, ACTINN) exhibited better performance, indicating the superior ability of artificial intelligence techniques in integrating large-scale data.

**Figure 4.**
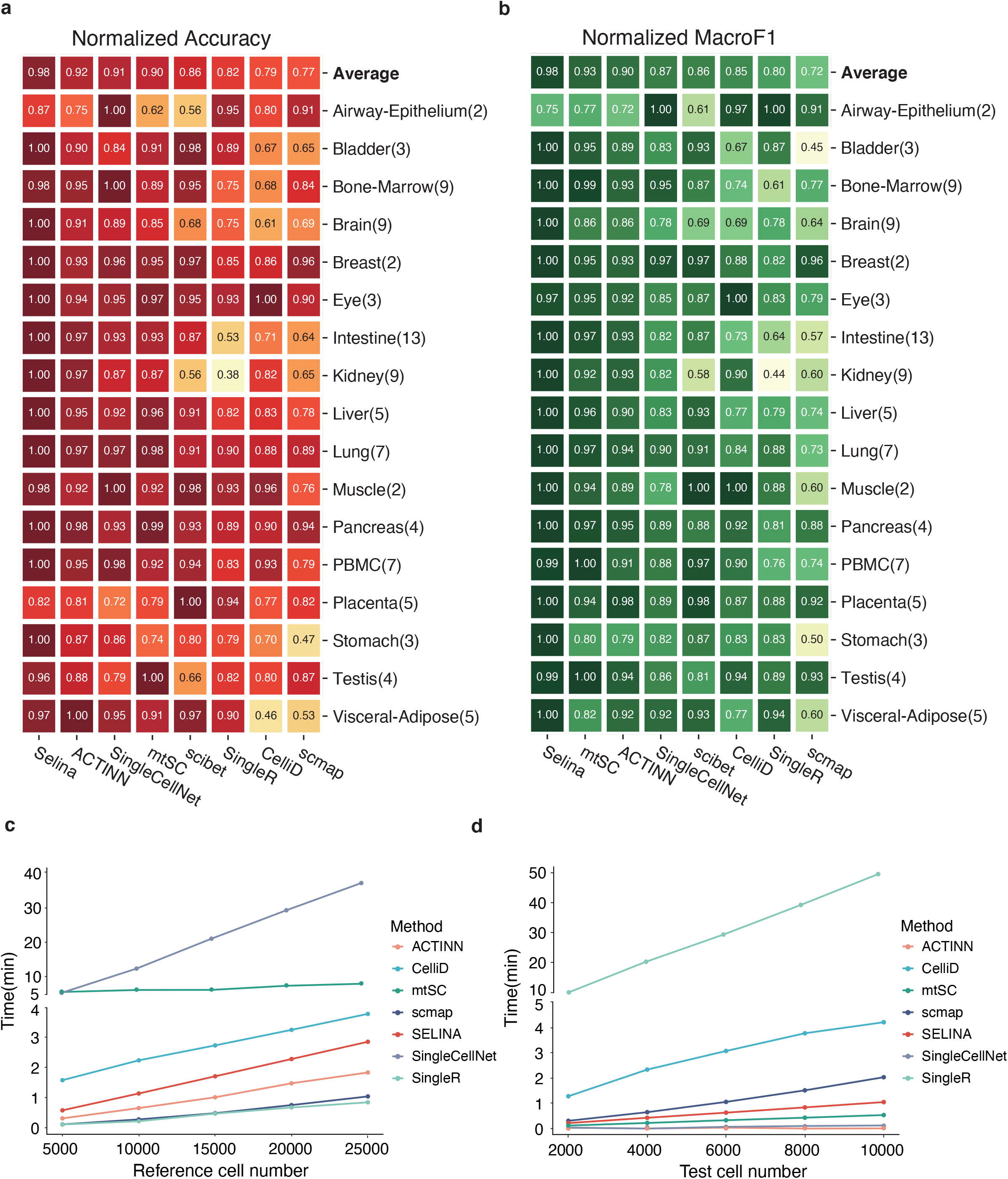
Performance evaluation of SELINA and existing annotation tools. **a.** The normalized accuracy for 17 tested tissues. The highest value in each tissue is regarded as 1, and other values are scaled according to the ratio to the highest value. On the top shows the average of all the tissues. **b.** The normalized MacroF1 for 17 tested tissues. Scales in the same way to the accuracy. **c.** Mean training time (3 repeats) for increasing training cell numbers. **d.** Mean testing time (3 repeats) for increasing testing cell numbers.

Moreover, we evaluated the training time and querying time of SELINA and the other tools. As SELINA, ACTINN and mtSC are deep learning-based methods, they were trained with a GPU and tested using a CPU, and the rest of the methods were trained and tested using the same CPU as the deep learning model testing used. We set various reference cell numbers and query cell numbers to investigate the dynamic change in consumption time and found that it was positively correlated with the cell number in all methods. Compared to other methods, SELINA exhibit a moderate computational efficiency. It takes SELINA approximately 3 minutes to train a model on a dataset with 25000 cells and less than 1 minute to fine-tune the parameters on a dataset with 10000 cells (Fig. 4c-d). Additionally, the fine-tuning step can be largely accelerated by the GPU with a more obvious effect as the cell number increases (Supplementary Fig. 4g). In conclusion, SELINA can realize more accurate annotation than published tools with an acceptable running time.

## Discussion

Substantial amounts of well-labeled human single-cell RNA-seq data have been generated in the public domain. Previous reference data-based studies utilized various strategies to achieve automatic annotation based on annotated reference scRNA-seq data. However, existing algorithms do not solve the problems of imbalanced cell types and large batch effects between reference and query datasets. In addition, due to the huge amount of public scRNA-seq data, a comprehensive and uniformed cell type reference is still not available. In this study, we developed an accurate deep learning-based framework SELINA for single-cell assignment along with a large-scale reference data portal covering 1,706,710 cells and 35 tissues. SELINA can handle the imbalance of cell types existing in reference data using SMOTE. In addition, SELINA can remove not only the batch effect across reference datasets but also the batch noise between the query dataset and the reference dataset. We systematically benchmarked the performance of SELINA on 17 different human tissues, and demonstrated its superiority in accurate cell type annotation compared to existing tools. Our method, combined with the curated reference, provides a one-stop solution for human single-cell annotations.

Conceptionally, SELINA splits the process of traditional transferring learning^15,18,26,27^ into two separate steps and consumes less time due to the separation of source data training and target data adaptation. The source data training is assigned to the software developer, and the target data adaptation is assigned to users. Thus, users can directly use the pretrained model in SELINA, perform an adaptive adjustment according to the query data, and accurately annotate the cell types within several minutes.

Despite the fact that our reference contains relatively large-scale datasets, the quantity of each dataset in different tissues is still imbalanced. This issue will be solved as the amount of data continues to grow or data generation algorithms^28,29^ develop. Additionally, different cells belonging to the same lineage with similar transcriptomic profiles are often misannotated in the reference data, e.g., CD4+ T cells and CD8+ T cells. Recently, CITE-seq^30^ and REAP-seq^31^ have been used to capture surface proteins that are helpful in distinguishing cells that are similar in transcriptomic profiles. By integrating the data from CITE-seq and REAP-seq, we can obtain more accurately annotated cell type references. Finally, the deep learning model we used only excels at clustering and classification tasks, it lacks biological insights, such as the ability to identify novel cell types or key factors. In the future, we will use graph-based algorithms to extract the structural information so that the association between genes and cell types can be preserved to present more explainable knowledge and used for novel cell type identifications. With the presence of the above features in the SELINA algorithm and continuous expansion of SELINA reference, we anticipate that SELINA to accurately characterize all human cell types with the potential to transfer to other species.

## Author contributions

C.W. and T.L. conceived the project. P.R. designed the SELINA algorithm and evaluated the performance. X.S., X.D., Z.Y., X.X.D., J.W., Y.Y., J.H. collected the data. X.S. processed the data with the help from P.R., Z.Y. and X.X.D., X.S. cleaned and built the SELINA reference. P.R., X.S., C.W. and T.L. wrote the manuscript with the help from other authors.

## Acknowledgements

This work was supported by the National Natural Science Foundation of China [32170660, 31801059, 81872290, 81972551, 81991502]. Shanghai Rising Star Program [21QA1408200]. Natural Science Foundation of Shanghai [21ZR1467600].

## Methods

### Reference construction

Based on single-cell relevant keywords, a text-mining-based data parsing procedure was created to collect published human disease-free single-cell RNA-seq data from several databases, such as GEO, EBI, HCA, and Broad, before 20210531. All of the datasets were double-checked to ensure that they met our requirements. Patient ID, tissue source, original cell type annotation, cell type markers, literature search number, sequencing platform, reference genome version, etc. were collected as well.

To make these datasets comparable, we created a standardized bioinformatics workflow. The data were preprocessed with MAESTRO^16^, which includes basic quality control, cell screening, normalization of expression data, unsupervised clustering, and differential expression analysis. Only cells with read count greater than 1000 and gene count greater than 500 were preserved. Batch effects from various patients or samples frequently affect datasets. To systematically measure the batch effect, each dataset was quantified with a metric based on the idea of information entropy and the Euclidean distance between cell coordinates in the UMAP graph. The batch effect was removed using the conventional correlation analysis^32^ (CCA) for data with entropy levels below the threshold.

The original cell type annotation taken from the papers was used to assure the authenticity and traceability of the data annotation. We used the cell type marker genes paper provided (Supplementary Table 3) to annotate datasets without original annotation from the corresponding papers. The average logFC of each cell type marker gene in each cell cluster was calculated as the cell type score, and the cell type with the highest score was assigned to that cell cluster. The annotation results were manually validated based on the expression distribution of the marker genes after the initial annotation.

The inconsistent annotations were first manually unified to the same label. Then we defined major lineage and minor lineage based on the cell types present in each tissue. For the minor lineage annotation unification, the cells assigned to the major lineage were divided into different minor lineages. As Supplementary Fig. 1b shows, cells belonging to endothelial lineage were extracted and we did PCA analysis, batch removal, re-clustering to these cells. Based on the marker genes papers provided, we re-annotated the endothelial cells into the sub-lineage cells. Cells without expression of any known marker genes were assigned with major lineage label attached with the suffix Subset. Finally, the cell type ontology tree was built for each tissue.

### Data augmentation of rare cell types

SMOTE is used to synthesize artificial samples of the minority class. A cell and the corresponding 5 nearest neighbors from the minority cell type are randomly chosen, and the artificially synthetic cell is generated at a random point on the line connecting the anchor cell and one of its selected neighbors. We denote the gene expression profiles of the anchor cell and neighboring cell as *x_a_* and *x_n_*, respectively, and the new cell *x_s_* is calculated using the following formula:

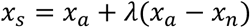

where *λ* is a random number between 0 and 1.

### Training and fine-tuning process of SELINA

The pretraining framework is composed of a feature extractor, cell type discriminator and platform discriminator. We denote the feature extractor as *G_f_* and the cell type discriminator as *G_c_*. Assuming we have *K* cell types, as the platform discriminator has an equal number of classifiers to cell types, these classifiers are denoted as 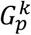, *k* = 1,…,*K*, each one is responsible for matching different platforms associated with one cell type. Suppose we have *N* cells; for each cell we have the gene expression vector *x_n_*, we denote the probabilities of one cell assigned to each of the K cell types as *ŷ_n_*, which is a vector calculated by

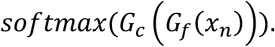

The attention of one certain platform discriminator paid to one cell was calculated by weighting the features with the corresponding probability in *ŷ_n_*. The cost function of the platform discriminator is calculated as the average of the loss of all classifiers within it, and the formula is shown as follows:

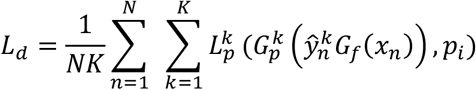

where 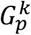 is the *k*-th platform discriminator with 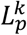 as its cross-entropy loss and *p_i_* is the platform label. The objective function is the sum of balanced loss from the cell type discriminator and platform discriminator, which is calculated using the formula listed below:

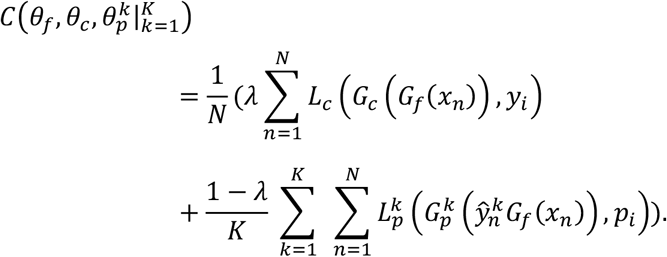

*L_c_* is the cross-entropy loss of the cell type discriminator, *y_i_* is the cell type label, and *λ* is a hyperparameter that balances the two objectives in the optimization problem. The optimization problem is to find the parameters 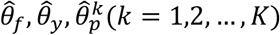, that simultaneously satisfy

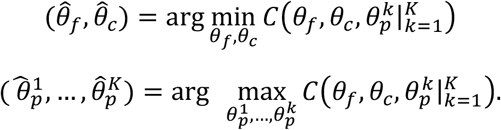

The feature extractor and cell type discriminator of the pretrained model are connected as the encoder of the autoencoder. We denote the encoder as *G_e_* and the decoder as *G_d_*. We first freeze the encoder and train the decoder and then fix the decoder and train the encoder. The objective function is

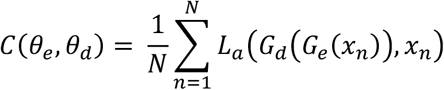

where *L_a_* is the mean squared error (MSE) loss of the autoencoder. The optimization problem is to find the parameters 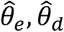 satisfying

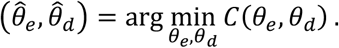

Once the fine-tuning step is finished, the decoder is removed, and the encoder is used to predict for unlabeled datasets.

### Model parameters

The neural network was implemented with PyTorch. The feature extractor contains three layers: one input layer with the same number of nodes as the gene number, one output layer with 100 nodes followed by a dropout layer. The cell type discriminator has four layers: a 100-node input layer, a 50-node hidden layer followed by a dropout layer, and an output layer with a number of nodes equal to cell types. Each platform discriminator unit contains three layers: a 100-node input layer, a 25-node hidden layer, and an output layer with a number of nodes equal to sequencing platforms.

The Rectified Linear Units (ReLU) was used as activation function. The Adam optimizer was used as the optimizer with default settings. The learning rates of pretraining and encoder training were set to 0.0001. For decoder training, the learning rate was set to 0.0005. The epoch numbers of pretraining and decoder training were set to 50, whereas the epoch number of encoder training was set to 20. The dropout layers’ parameters were set to default.

### Benchmark of SELINA and existing tools

For all datasets in each tissue, we iteratively took one dataset as a query and merged the remaining datasets as a reference dataset. The extremely large datasets that consume vast amounts of memory were downsampled with all cell types intact. The expression profiles of the reference data and query data were scaled to 10000 and log-transformed. The querying process used common genes between query and reference data.

For all benchmarks, scmap-cell, singleR, SingleCellNet, scibet and CelliD were trained and tested with CPU AMD EPYC 7552 2.2 GHz. SELINA, ACTINN, and mtSC, which are deep learning-based frameworks, were trained with GPU GTX960 and tested with AMD EPYC 7552 2.2 GHz. All the parameters were the defaults or set as recommended in the corresponding documentations.

### Evaluation metrics

The accuracy was defined as the ratio of corrected assigned cells over all cells. The MacroF1 score was calculated as listed below:

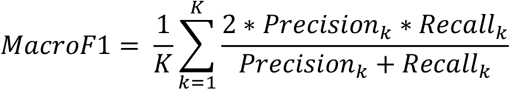

*K* represents the number of cell types in the query dataset. *Recall_k_* and *Recall_k_* are the recall and precision of the *k*-th cell type.

### Data and codes availability

For added convenience, we provide both the SELINA Python package and the pretrained models covering 134 datasets (Supplementary Table S1). The source code of SELINA is deposited on GitHub (https://github.com/wanglabtongji/SELINA) and can be easily installed via conda. The pretrained models are available from (https://github.com/wanglabtongji/SELINA_reference)

## Supplementary information

**Supplementary Figure 1 Pipeline for data processing and cell type curation.**

**a.** The processing procedure for the individual data. First the low quality cells are filtered and the dimension of the expression profiles from the remaining cells are reduced with PCA. The batch effect across different samples or patients within a dataset is removed. Then the cells are clustered and assigned with cell types based on either the original annotation or the marker genes papers provided.

**b.** Annotation harmonization across datasets in each tissue. Here we take the endothelial cells as the example. Cells belonging to endothelial lineage are extracted. Then PCA analysis, batch removal and re-clustering are applied. Based on the marker genes papers provided, the endothelial cells are re-annotated into its sublineage cells. Cells without expression of any known marker genes are assigned with major lineage label attached with the suffix Subset.

**Supplementary Figure 2 Data summary of the reference data atlas.**

Cell number (**a**) and cell type number (**b**) for each tissue.

**Supplementary Figure 3 Validation of the strategies used in SELINA.**

**a-b.** Low-dimensional representation of the reference data from liver after feature extractor transformation (**a**) and cell type classifier transformation (**b**).

**c.** Mean accuracy (3 repeats) of different strategies. Base represents the situation in which the data is not oversampled and the pre-trained model is used without finetuning. SMOTE represents the results when SMOTE is additionally implemented to oversample rare cell types. SMOTE+Autoencoder shows the results when SMOTE and autoencoder are both implemented.

**d.** Performance test of multiple datasets and a single dataset used as reference data in lung.

**Supplementary Figure 4 Performance evaluation of SELINA and existing annotation tools.**

**a-e.** Performance of tested tools on tissues with a large number of datasets.

**f.** Benchmark for the time consumption using CPU and GPU in fine-tuning step. Each point represents the mean value of three replicates.

**Supplementary Table 1 Datasets collected in the SELINA reference atlas**

**Supplementary Table 2 Datasets used in the benchmark of SELINA and existing annotation tools**

**Supplementary Table 3 Marker genes from the original papers**

