## Supplementary figures and images for "SELINA: Single-cell Assignment using Multiple-Adversarial Domain Adaptation Network with Large-scale References"

### Supplementary Figures

#### Supplementary  Figure 1

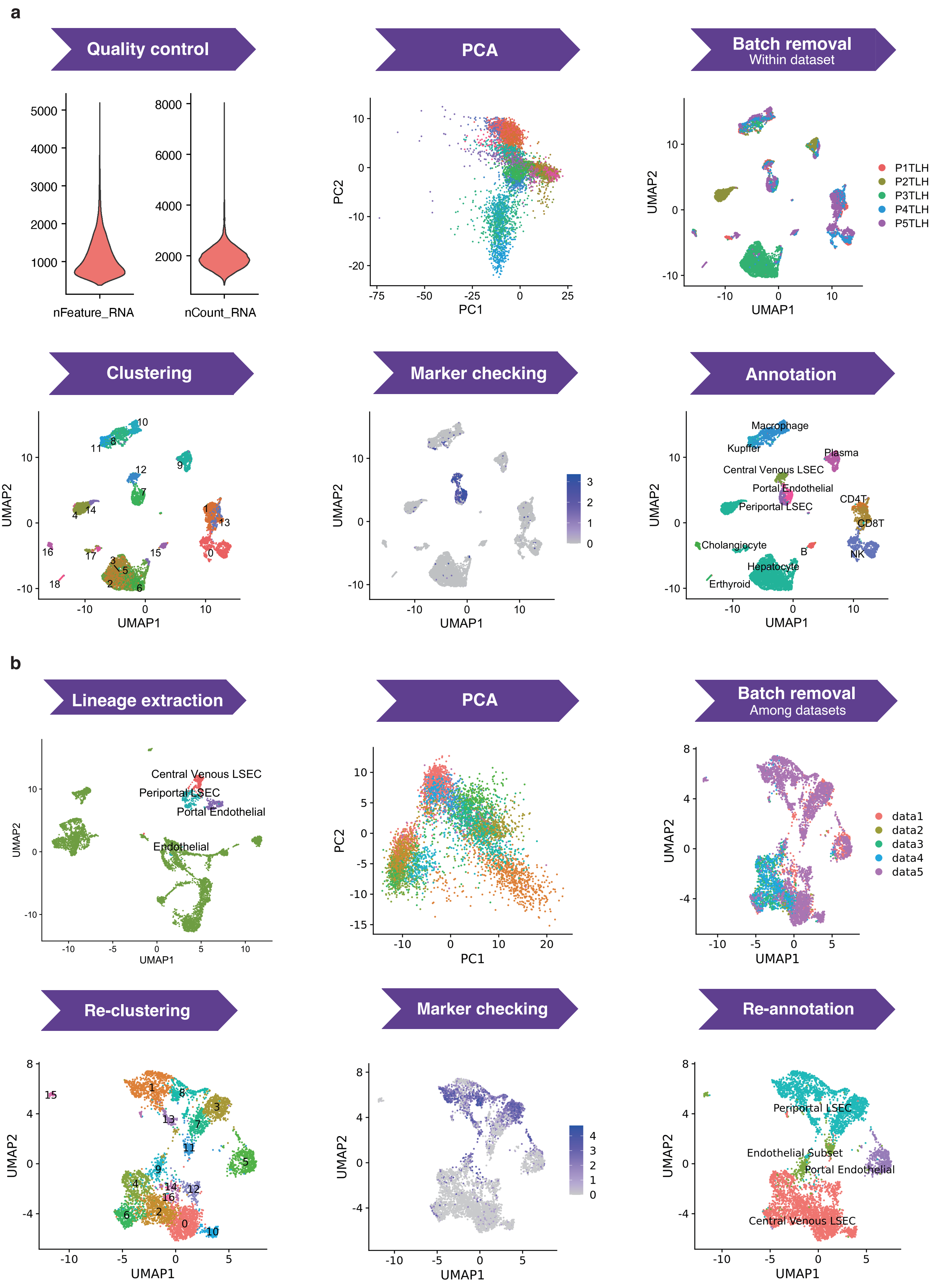

#### Supplementary Figure 2

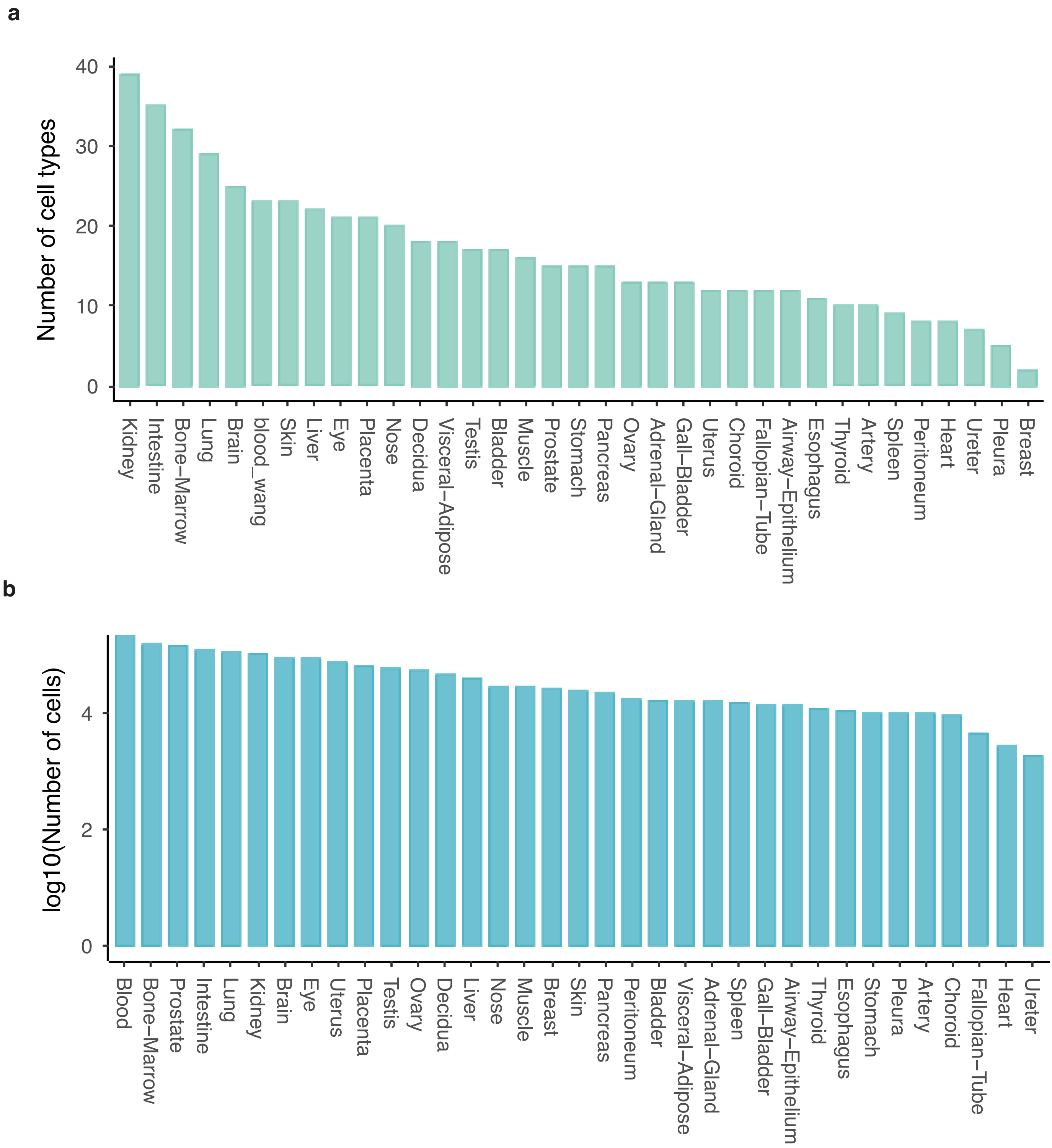

#### Supplementary Figure 3

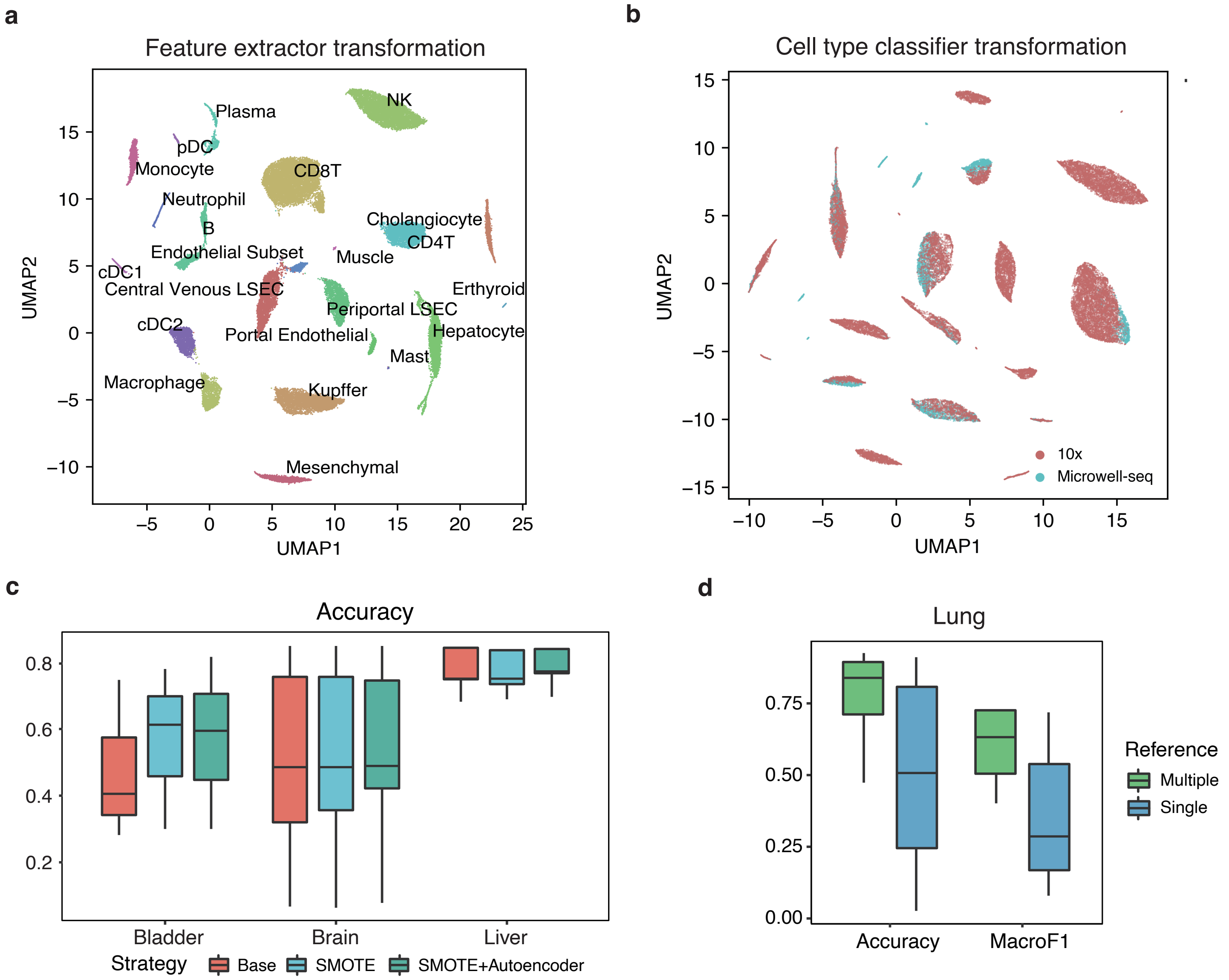

#### Supplementary Figure 4

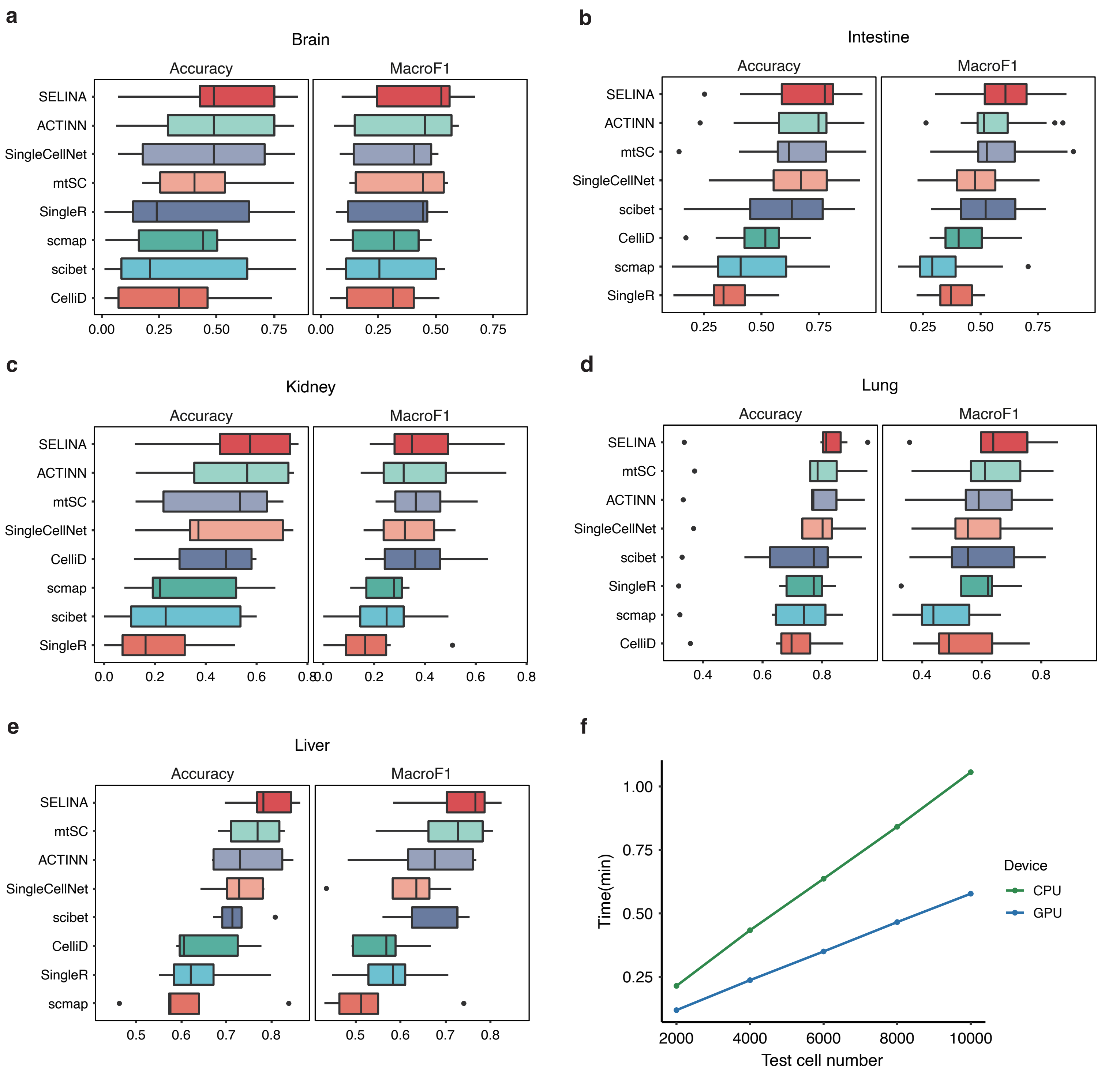
